# Time- and Dose-dependent Effects of CIGB-300 in the lung squamous cell carcinoma proteomics

**DOI:** 10.1101/2024.12.03.626528

**Authors:** Liudy García-Hernández, Lingfeng Dai, Arielis Rodríguez-Ulloa, Ying Yi, Luis J. González, Vladimir Besada, Wen Li, Silvio E Perea, Yasser Perera

**Author notes:** Correspondence (Y.P); (S.E.P.); (W.L). Equal Contribution (Liudy García-Hernández, Lingfeng Dai). (L.G), (A.R.U.); (L.J.G.); (V.B.), (Y.P); (L.D); (Y.Y); (W.L), Cuba; (S.E.P.).

## Abstract

Proteome-wide scale in a dose- and time-depending setting is crucial in drug-regulated proteomics to fully understand the pharmacological mechanism of anticancer drugs as well as the identification of proteins suitable for drug response biomarkers. Here, we investigated the effect of the CIGB-300 anticancer peptide at IC_50_ and IC_80_ dose levels during 1 and 4 hours of treatment on squamous lung cancer cell (NCI-H226) proteome. A dose-dependent inhibitory effect of the global proteomics while slight change of the upregulated proteins was observed, by increasing CIGB-300 dose level. The biological processes that were more represented among the modulated proteins were those related to small molecule biosynthetic process, pyridine-containing compound metabolic process and nucleobase-containing small molecule metabolism. Importantly, NSCLC previously described biomarkers and Protein Kinase CK2 substrates were significantly modulated by treating both doses of CIGB-300. These proteins, which are associated with both metabolism and cell survival, reinforce the hypothesis of the therapeutic potential of CIGB-300 as anticancer treatment for squamous non-small cell lung cancer and merits be validated as CIGB-300 response biomarkers in NSCLC patients. Overall, our proteomics-guided strategy based on time and drug dose served not only to deep into the mechanism of action of CIGB-300 but also to identify candidate proteins for pharmacodynamic and drug response biomarkers.

## Introduction

Lung cancer is the most common oncologic disease worldwide and the leading cause of cancer-related mortality (1.8 million deaths, 18.7% of total cancer deaths in 2022) (Bray et al. 2024). It has traditionally been classified into two primary groups; small cell lung cancer and non-small cell lung cancer (NSCLC). Lung adenocarcinoma (LUAD) and lung squamous cell carcinoma (LSCC) are the most common subtypes of the last one (Herbst et al. 2008). NSCLC represents approximately 85% of these, is associated with smoking and causes approximately 40,000 to 50,000 deaths annually in the United States (Liao et al. 2012).

Several therapies have been developed and applied to treat this disease, and significant progress has been made in this field. However, it still remains an unmet scientific challenge. In particular, patients with lung squamous cell carcinoma (LSCC), unlike lung adenocarcinoma, have not benefited from effective or targeted therapies (Niu et al. 2022).

Casein kinase 2 (CK2) is involved in several metabolic and regulatory processes in cells, accounting for approximately 24.1% of the cellular phosphoproteome. These processes include apoptosis or programmed cell death and cell proliferation. It is also involved in DNA repair mechanisms, which has been linked to resistance to antitumor drugs. Upregulation of CK2 in cells is an indicator of cancer and has been associated with worse prognosis; which supports its considerable relevance as a target for study and development of possible treatments against tumor phenotype (Borgo et al. 2021).

CIGB-300 is a synthetic anti-cancer peptide developed by Perea and colls. (Perea et al. 2018). It contains a cyclic peptide that inhibits (CK2) activity in cancer cells through peptide-substrate interaction, preventing phosphorylation and thus its regulatory activity (Perea et al., 2018, 2004). Recently, it has been demonstrated by cell-free assay that CIGB-300 directly interacts with the CK2 catalytic subunit alpha and consequently blocks the kinase activity, which is a parallel mechanism to the one previously described and complementary to the anticancer effect (Perera et al. 2020).

The biological effects and non-clinical antitumor activity of CIGB-300 have been previously studied in various types of cancer, including SCLC, NSCLC and cervical cancer (Perea et al. 2008; Perera et al., 2009, 2015)). This peptide is also able to inhibit the proliferation, adhesion and invasion of lung cancer cells by interfering with key signaling pathways associated with malignant progression ((Perea et al. 2008; Perera et al., 2009). In addition, non-clinical studies in cervical and lung cancer models have shown that systemic administration of CIGB-300 in syngeneic mice is associated with a significant reduction in tumor growth and angiogenesis, as well as inhibition of cancer cell dissemination and colonization of distant organs (Perea et al. 2018). Also, a Phase 1 study in patients with advanced solid tumors showed that lung cancer patients tended to exhibit longer survival time after CIGB-300 treatment (Batista-Albuerne et al. 2018).

Proteomic analysis can be a powerful tool to obtain direct evidences of CIGB-300 effect and mechanism of action in squamous NSCLC, metabolic pathways and cellular processes impaired by this inhibitor. In fact, previous proteomics data on CIGB-300-treated NSCLCs revealed different biological processes and pathways involved on the drug action (Rodríguez-Ulloa et al. 2010). However, there is a lack of dose- and time-dependent CIGB-300 characterization at the level of proteins. To address this gap, here we presents a quantitative proteomic approach uncovering proteins engaged in the CIGB-300 mechanism in a dose- and time-resolved fashion by using label-free shotgun proteomic analysis. Particularly, we interrogated the temporal cellular proteome that is regulated at two different pharmacological perturbating dose levels, the Inhibitory Concentration -50 (IC_50_) and IC_80_.

## Material and Methods

### Cell culture and sample preparation

NCI-H226 (10^7^ cells per each condition, three biological replicates) were incubated with IC_50_ (113µM) and IC_80_ (135.6 µM) of CIGB-300 peptide during 1h and 4 h. Cells were washed three times with 10 mL PBS and centrifuged for 5 min 1000 g. Cell pellets were resuspended in 200 μL of sample buffer (0.1 M Tris HCL, 0.05 M DTT, 1.5 % SDS pH 8.0) for lysis by heat at 95°C for 5 min and sonicated for 3 cycles of 15 seconds each one in an ultrasonic processor (Model GE 50 T 20 kHz, Markson). Protease and phosphatase inhibitor cocktails Complete TM and PhoStop TM (Roche Diagnostics) respectively were added to the resuspended pellets. The lysates samples were centrifuged at 18,000 rpm.g for 20 min, 4 °C. For all the samples, precipitate was discarded and the supernatant was then collected for the proteome analysis. Afterwards the samples were diluted and total protein concentration was measured using the X protein assay kit. Aliquots containing 200 μg of proteins were subjected to digestion with trypsin by filter-aided sample preparation as described by Wisniewski et al (Wiśniewski et al. 2009).

### NanoLC-MS/MS and Data Analysis

For LC-MS/MS runs, the tryptic peptides mixtures corresponding to 1μg proteins were injected via nanoLC Ultima 3000 coupled with a Pepmap column (75 μm x 150 mm) to the Thermo Exploris 480 mass spectrometer (with FAIMS). Gradients of 95% solution B (acetonitrile 80% in 0.1% formic acid) wereperformed in 65 minutes at 300 nL/min flow rate. The mass spectrometer was operated in data dependent analysis (DDA) mode with dynamic exclusion of 30 s and full-scan MS spectra (m/z 350–1500) with resolution of 120 000, followed by fragmentation of the most intense ions within 1 s cycle time with high energy collisional dissociation (HCD), normalized collision energy (NCE) of 30.0 and resolution of 15,000 (m/z 200) in MS/MS scans. Peptide/protein identification was performed by Proteome Discoverer v.2.4.0.287 (Thermo Scientific, USA) using the “uniprotkb_HUMAN_20240115.fasta” defining as fixed modification the cysteine carbamidomethylation and as dynamic modification the deamidation (N/Q) and methionine oxidation. Trypsin was the enzyme selected for the digestion, allowing up to one missed cleavage sites per peptide. A false discovery rate of 0.05 was considered for identification.

### Bioinformatic Analysis

Biological processes, biological pathways and oncogenic signatures significantly represented in differentially modulated proteins were identified through functional annotation and enrichment analysis, based on the information annotated in the Gene Ontology (GO, http://www.geneontology.org), KEGG PATHWAY (https://www.genome.jp/kegg/), Reactome (https://reactome.org/) and MSigDB (https://www.gsea-msigdb.org/gsea/msigdb/index.jsp) databases (Ashburner et al. 2000; Gillespie et al. 2022; Kanehisa et al. 2000; Liberzon et al. 2011, 2015). Analysis was performed with Metascape gene annotation and analysis resource (https://metascape.org/), a web-based tool that computes the accumulative hypergeometric distribution and enrichment factors to identify significantly enriched biological processes through statistical analysis (p-value <0.01, enrichment factor >1.5) (Zhou et al. 2019). For the proteomic profile regulated at 113µM (IC50) and 135.6 µM (IC80) of the CIGB-300 peptide, the Metascape Custom Analysis option was selected [accessed on October 28-29, 2024], the NCI-H226 cell line proteome was used as background, and the modulated proteins at 1h and 4h were used as independent input data sets for a time-dependent meta-analysis.

CK2 substrates differentially modulated in NCI-H226 cells after CIGB-300 treatment were retrieved using the PhosphoSitePlus database (http://www.phosphosite.org (accessed on October 29, 2024)) (Hornbeck et al. 2019). The STRING database (http://string-db.org/ (accessed on November 4, 2024) (Szklarczyk et al. 2019) was used to identify interactions between differentially modulated proteins. In addition to the CIGB-300 regulated proteome, proteins associated with oncogenic signatures over-represented in the proteomic profile were also included for protein interaction network analysis. In such analysis only text mining, databases and experimental evidences were used as sources of interaction data and the confidence score was fixed at 0.7. Biological processes and protein complexes represented in protein-protein interaction networks were retrieved by STRING functional enrichment tool. Protein-protein interaction networks were visualized using Cytoscape software (v.3.9) (Shannon et al. 2003).

## Results

### Temporal proteomic profile of the NCI-H226 cell line treated with the CIGB-300 at two dose levels

Firstly, the temporal proteomic profile of NCI-H226 cells treated with CIGB-300 at IC_50_ and IC_80_ during 1 and 4 hours was identified by using shotgun proteomics and Proteome Discoverer label-free quantitative analysis (Table S1). A total of 4700 proteins were detected for the cells treated with IC_50_ and IC_80_ CIGB-300 at 1h incubation time. After applying statistical analysis and normalization to the total number of identified proteins (p<0.05 and ≥ 2 fold-change), it was found that 427 were differentially modulated by CIGB-300 at IC_50_ dose (122 up-regulated and 305 down-regulated) and 720 proteins at IC_80_ dose (143 up-regulated and 577 down-regulated) after 1h of treatment (Fig. 1). After 4h of incubation, 4510 proteins were identified and 351 were differentially modulated by the peptide treatment. Among regulated proteins, 64 were up-regulated and 287 down-regulated at IC_50_ dose, while there was also a significant change in the protein abundance for 571 of them with IC_80_ (78 up-regulated and 493 down-regulated) (Fig. 1). In all the four experimental conditions, the numbers of CIGB-300 down-regulated proteins were higher than those upregulated by the peptide (Fig. 1). Interestingly, the CIGB-300 treatment elicited a dose-dependent downregulation of the global NCI-H226 cellular proteome both after 1 and 4 hours of incubation. Otherwise, the number of proteins that were up-regulated was nearly unchanged by increasing the CIGB-300 dose. Of note, the temporal dynamic of down- and up-regulated proteins seemed to be unchanged at CIGB-300 IC_50_ dose while slight time-dependent decay was observed at IC_80_ peptide dose. (**Error! Reference source not found.**). Thus, the CIGB-300 regulated proteome seems to be scarcely modified between 1 and 4 hours.

**Figure 1.**
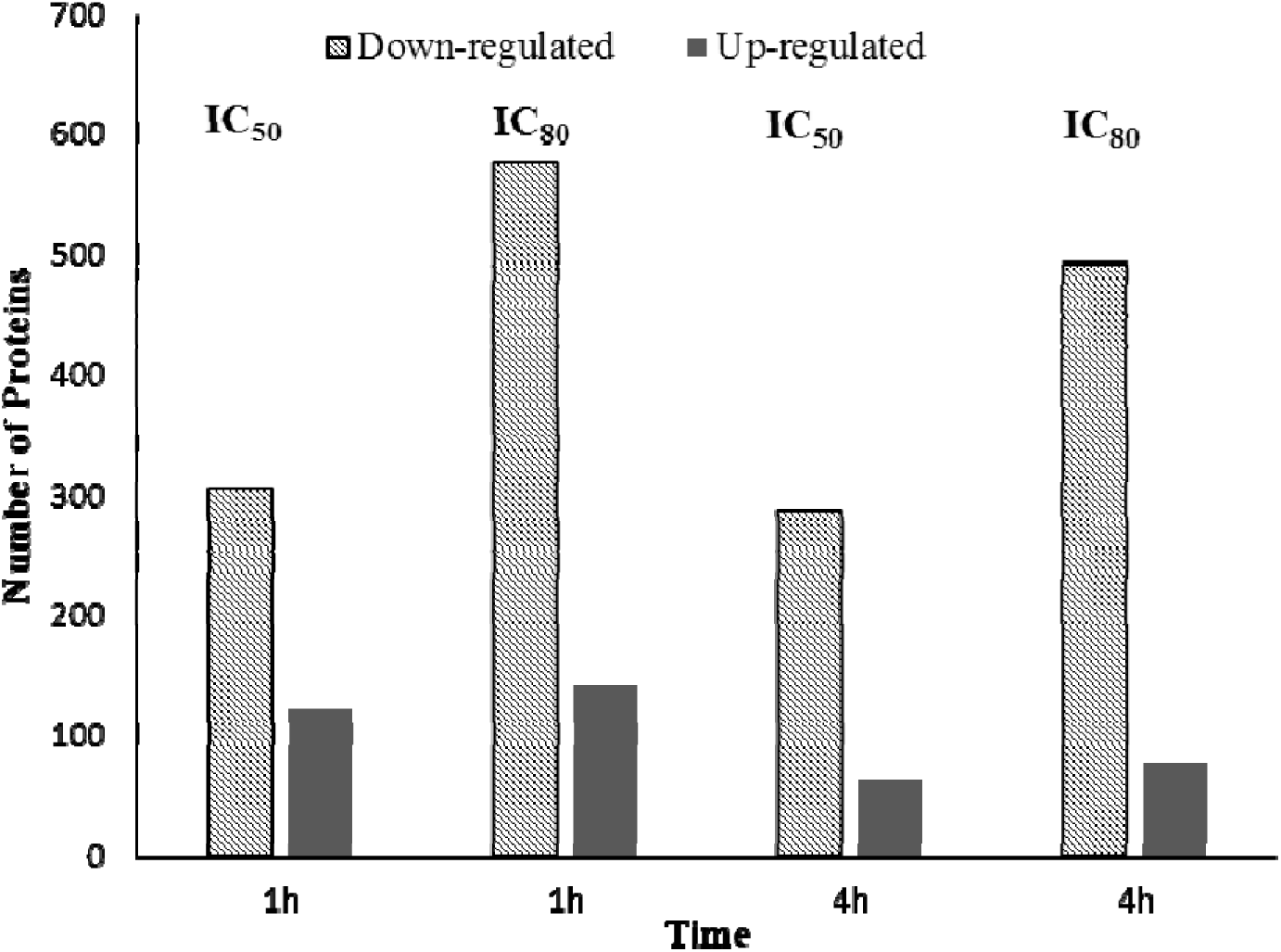
Differentially expressed proteins (FC≥2, p < 0.05) in NCI-H226 cell line treated with CIGB-300 for IC50 and IC80 doses during 1h and 4 h.

### Functional Characterization of CIGB-300 Regulated Proteome in NCI-H226 cells

Functional enrichment analysis of the CIGB-300-regulated proteome in NCI-H226 cells was performed using the Metascape bioinformatic tool [**Error! Bookmark not defined.**]. In such analysis, proteins modulated by CIGB-300 IC_50_ treatment were related to biological processe and pathways that were also over-represented in the proteomic profile of NCI-H226 cells treated with CIGB-300 IC_80_ (Fig. 2) (Table S1). In contrast to the proteome regulated by CIGB-300 IC_50_ and CIGB-300 IC_80_ at 1h, a wider range of functional annotations merged between the proteomic profiles modulated after 4h of treatment with both CIGB-300 doses. In general, these annotations have a higher enrichment *p*-value in the proteome regulated by CIGB-300 IC_80_ (Fig. 2). Proteins involved in small molecule biosynthesis, aldehyde metabolism and metabolism of nucleobase / pyridine – containing compounds were modulated in CIGB-300-treated cells irrespectively to the peptide dose or the treatment time (Fig. 2). The proteome modulated by both CIGB-300 doses after 4h is characterized by an over-representation of proteins related to metabolic processes that supports the proliferative capacity of NCI-H226 cells, such as the pentose phosphate pathway and the metabolism of amino acids, fructose and mannose (Fig. 2 B). In addition, CIGB-300 modulates proteins involve in the endosomal and vacuolar pathway, this functional category includes a subset of proteins implicated with the immune cell mediated cytotoxicity.

**Figure 2.**
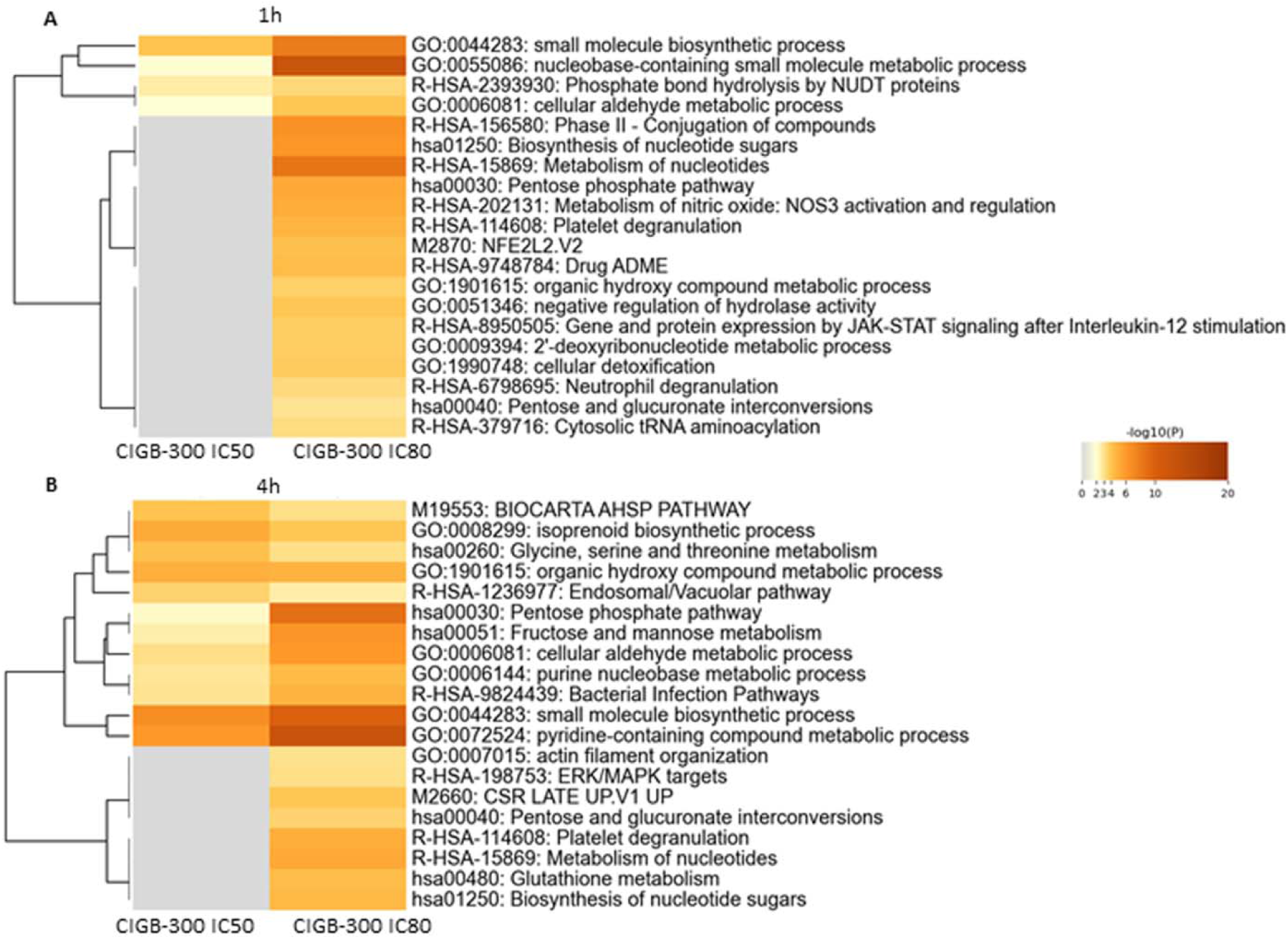
Enrichment analysis for differentially modulated proteins in NCI-H226 cells treated with CIGB-300 IC_50_ (113 µM) or CIGB-300 IC_80_ (135.6 µM) during: (A) 1h and (B) 4h. Biological processes and pathways significantly represented in the proteomic profiles (*p*-value < 0.01, enrichment factor > 1.5), were identified using Metascape gene annotation and analysis resource (https://metascape.org/). In the heatmap enriched terms are colored according to *p*-values.

Several biological processes and pathways are over-represented only in the proteomic profile regulated by CIGB-300 IC_80_, illustrating a dose-dependent effect. The proteomic profile differentially modulated by CIGB-300 IC_80_ at 1h includes proteins of the oncogenic signature NFE2L2.V2 (Fig. 2 A). Proteins encoded by genes that are down-regulated following the inhibition of NFE2L2/NRF2 (Kim et al. 2016), were found to be down-regulated in response to CIGB-300 IC_80_ treatment. Importantly, CK2 regulates the NRF2-mediated cellular antioxidant response (Pi et al. 2007). After increasing the concentration of CIGB-300 up to IC_80_, JAK-STAT and ERK/ MAPK mediated signaling events were modulated in NCI-H226 cells at 1h and 4h, respectively (Fig. 2). Both signaling cascades are activated by CK2 (Plotnikov et al. 2019; Zheng et al. 2011) therefore modulation of such pathways could be a direct down-stream effect of the CK2 inhibition mediated by CIGB-300 IC_80_.

The cellular response to a higher cytotoxic concentration of CIGB-300 is characterized by an over-representation of proteins related to cellular detoxification and glutathione metabolism which were modulated at 1h and 4h after treatment with CIGB-300 IC_80_. Concomitantly, proteins related to drug ADME (Absorption, Distribution, Metabolism and Excretion) properties were over-represented in the proteomic profile modulated at 1h (Fig. 2A).

### Network analysis of CK2 substrates modulated by CIGB-300

To further characterize the proteomic profile regulated by CIGB-300, the CK2 substrates differentially modulated in the peptide–treated NCI-H226 cells were retrieved from the Phosphosite Plus database (Hornbeck et al. 2019). A total of 15 CK2 substrates were differentially modulated in response to CIGB-300 IC_50_ treatment, with the majority (9/15) showing decreased abundance upon CK2 inhibition by peptide (Fig 3 A). In the presence of CIGB-300 IC_80_ a higher number of CK2 substrates were modulated, four decreased and 15 increased their abundance levels in peptide-treated NCI-H226 cells (Fig 3 B). A subset of ten CK2 substrates were equally modulated at both CIGB-300 doses (Fig 3).

**Figure 3.**
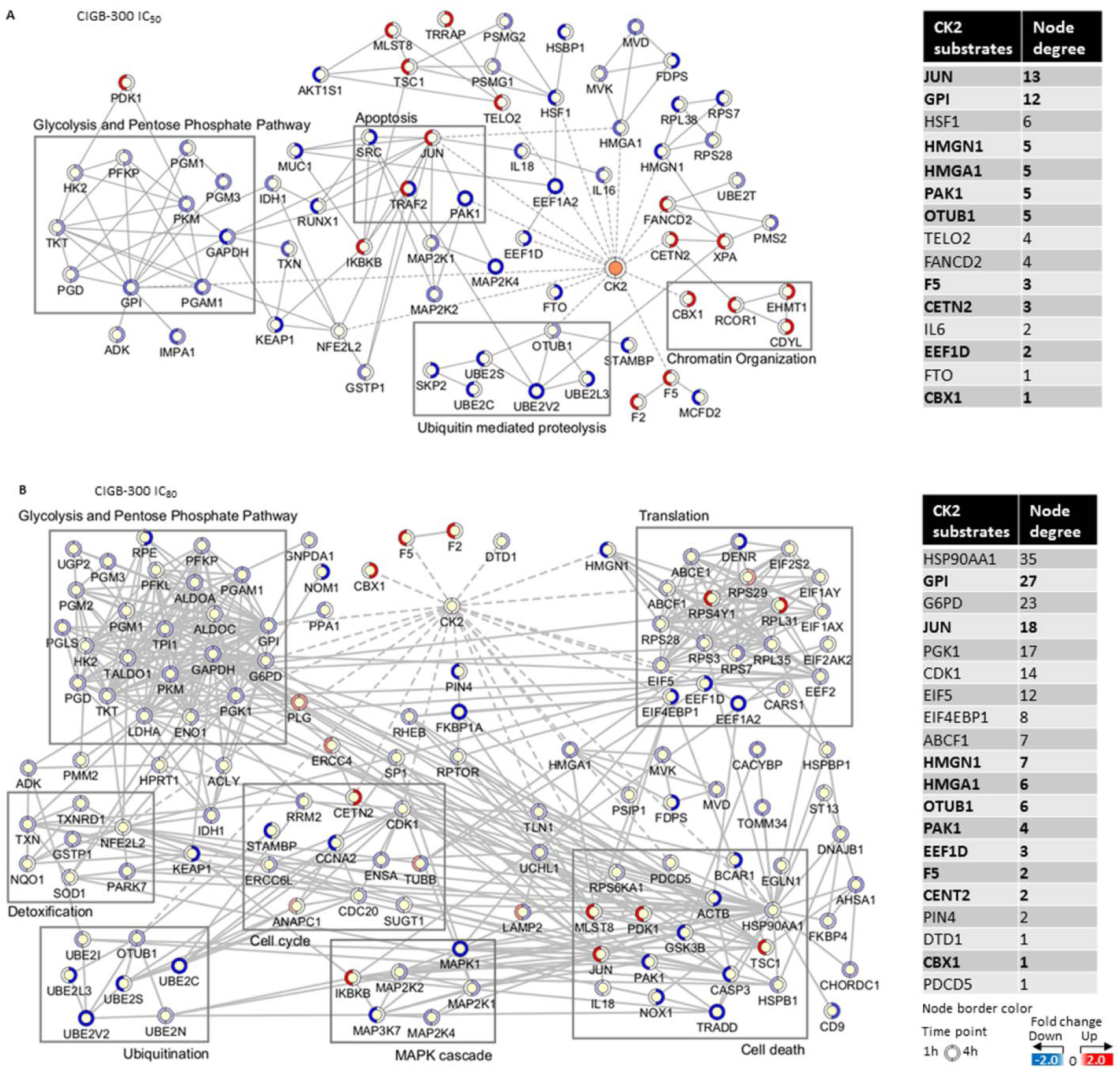
Protein-protein interaction networks associated with the proteomic profile modulated by CIGB-300 IC_50_ (A) and CIGB-300 IC_80_ (B) in NCI-H226 cells. In both networks, proteins are represented according to their expression level at 1h and 4h in a clockwise fashion (blue, decrease; red, increase; yellow, not identified). Biological processes and protein complexes gathered using the STRING functional enrichment tool are indicated by squares. Interactions between CK2 and its substrates are represented by dashed edges. For each CK2 substrate, the number of interactions (degree) is indicated in the tables, with those differentially modulated by CIGB-300 IC_50_ and CIGB-300 IC_80_ highlighted in bold.

The interaction network among the differentially modulated proteins was represented using the STRING database (Szklarczyk et al. 2019). The interactome around the CK2 substrates modulated in response to peptide treatment contains a total of 133 and 506 direct interactions connecting 55 and 112 proteins differentially modulated at CIGB-300 IC_50_ and CIGB-300 IC_80_, respectively. CIGB-300 IC_50_ modulated CK2 substrates with the highest number of interactions include the transcription factors JUN and HSF1, and the glycolytic enzyme GPI (Fig 3). In the case of the CIGB-300 IC_80_ regulated proteome, the CK2 substrates with the highest number of network connections include the protein kinases PGK1 and CDK1, and the chaperone protein HSP90AA1 (Fig 3). Such proteins, which have a high degree of network interactions, might exert transcriptional and post-translational regulatory functions downstream of CK2 inhibition by CIGB-300, supporting its mechanism of action.

As illustrated in Figure 3, CK2 substrates down-regulated by CIGB-300 IC_50_ and IC_80_ are located in clusters of proteins related to ubiquitin-mediated proteolysis, glycolysis and pentose phosphate pathway. Consistent with the pro-apoptotic effect of CIGB-300 in lung cancer cells (Perea et al. 2008), cell death related proteins were differentially modulated at both concentrations of the peptide, reflecting an impairment of the CK2 regulated cell survival events (Fig 3). On the other hand, clusters of proteins related to translation, cell cycle, MAPK cascade and detoxification were modulated in response to CIGB-300 IC_80_. In the proteome modulated by the lower dose of CIGB-300, such downstream effects of CK2 inhibition were barely represented.

### CIGB 300 inhibit key proteins upregulated in Non–Small Cell Lung Cancer and squamous NSCLC

In this work, label-free shotgun proteomic analysis permitted to detect low abundance proteins and provide quantitative information. Data from Table 1 show several key proteins or potential biomarker usually up-regulated in NSCLC and squamous NSCLC that were inhibited by CIGB-300 in this study (Table 1).

**Table 1.**
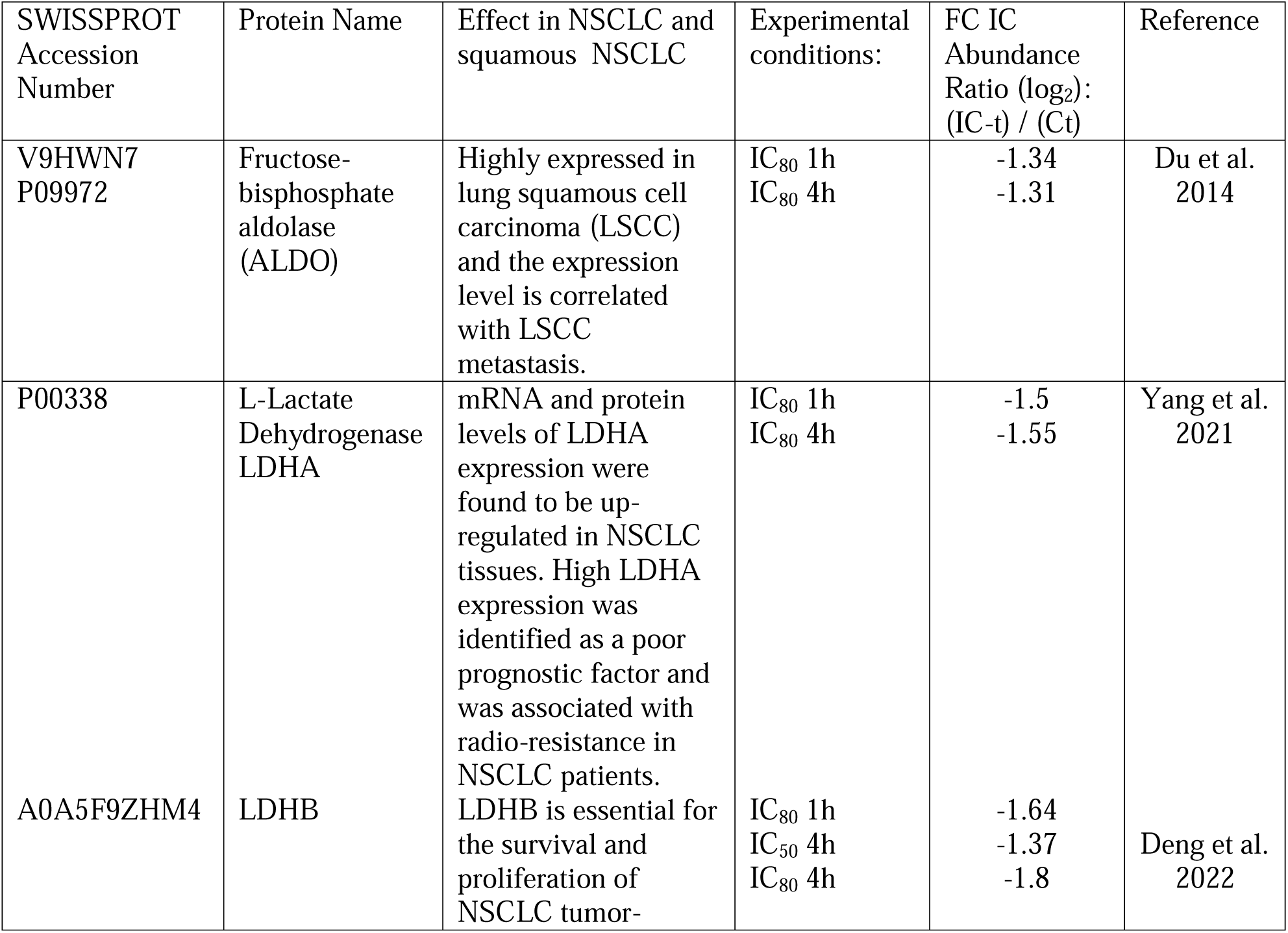
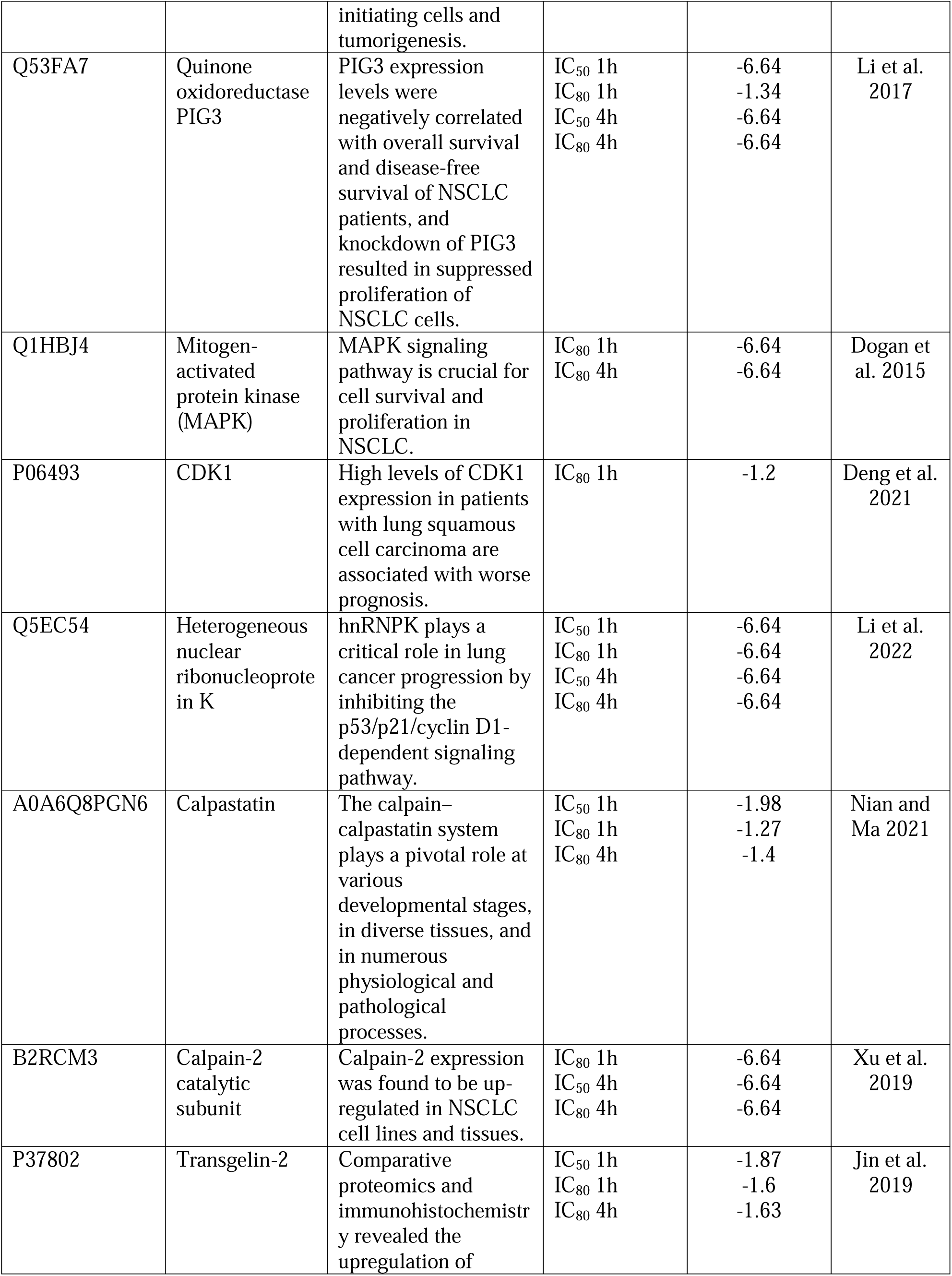
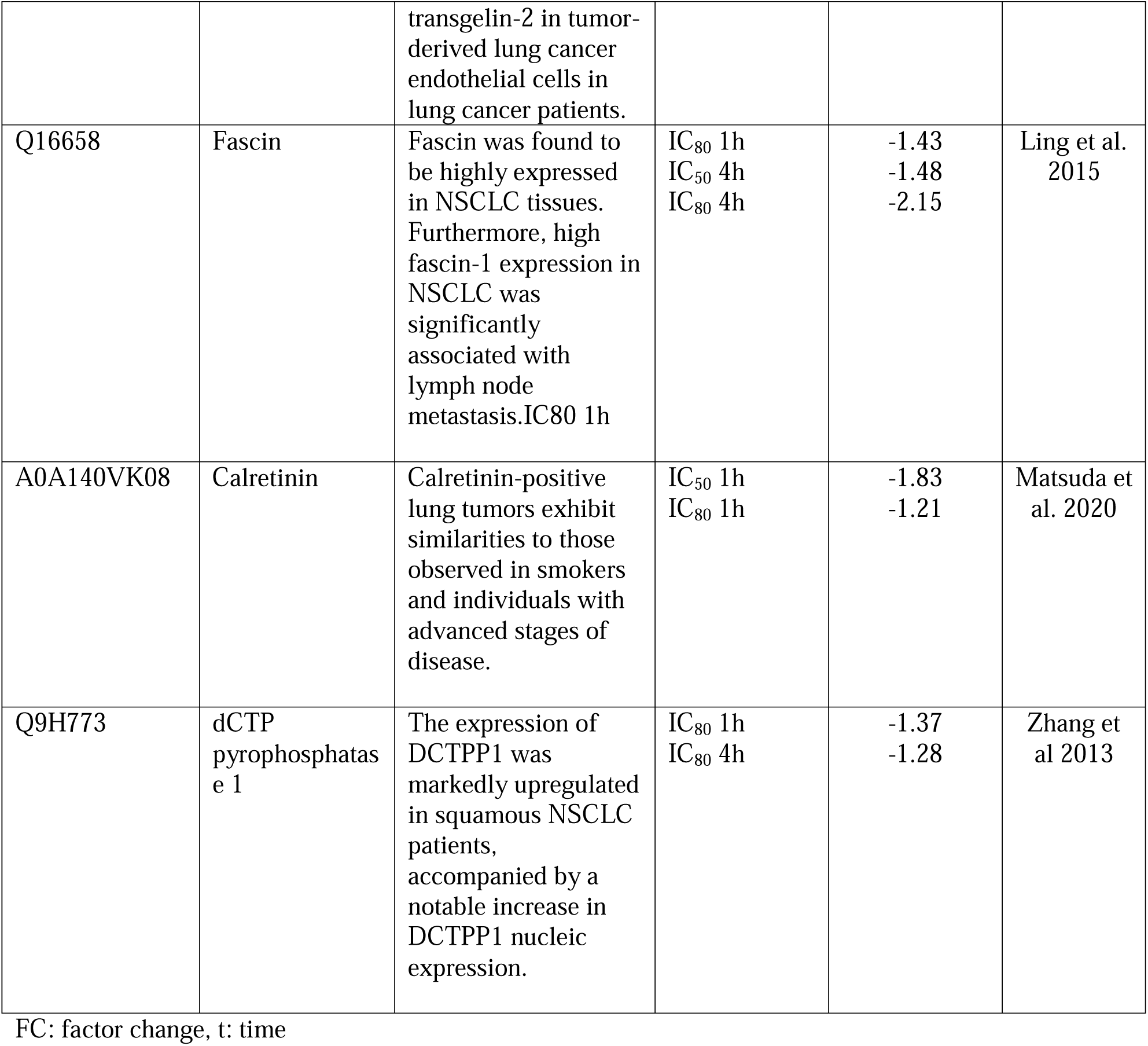
Down-regulation of NSCLC-associated proteins by CIGB-300 on NCI-H226 cells.

| SWISSPROT Accession Number | Protein Name | Effect in NSCLC and squamous NSCLC | Experimental conditions: | FC IC Abundance Ratio (log<sub>2</sub>): (IC-t) / (Ct) | Reference |
| --- | --- | --- | --- | --- | --- |
| V9HWN7<br>P09972 | Fructose-bisphosphate aldolase (ALDO) | Highly expressed in lung squamous cell carcinoma (LSCC) and the expression level is correlated with LSCC metastasis. | IC<sub>80</sub> 1h<br>IC<sub>80</sub> 4h | -1.34<br>-1.31 | Du et al. 2014 |
| P00338 | L-Lactate Dehydrogenase LDHA | mRNA and protein levels of LDHA expression were found to be up-regulated in NSCLC tissues. High LDHA expression was identified as a poor prognostic factor and was associated with radio-resistance in NSCLC patients. | IC<sub>80</sub> 1h<br>IC<sub>80</sub> 4h | -1.5<br>-1.55 | Yang et al. 2021 |
| A0A5F9ZHM4 | LDHB | LDHB is essential for the survival and proliferation of NSCLC tumor- initiating cells and tumorigenesis. | IC<sub>80</sub> 1h<br>IC<sub>50</sub> 4h<br>IC<sub>80</sub> 4h | -1.64<br>-1.37<br>-1.8 | Deng et al. 2022 |
| Q53FA7 | Quinone oxidoreductase<br>PIG3 | PIG3 expression levels were negatively correlated with overall survival and disease-free survival of NSCLC patients, and knockdown of PIG3 resulted in suppressed proliferation of NSCLC cells. | IC<sub>50</sub> 1h<br>IC<sub>80</sub> 1h<br>IC<sub>50</sub> 4h<br>IC<sub>80</sub> 4h | -6.64<br>-1.34<br>-6.64<br>-6.64 | Li et al.<br>2017 |
| Q1HBJ4 | Mitogen-activated protein kinase (MAPK) | MAPK signaling pathway is crucial for cell survival and proliferation in NSCLC. | IC<sub>80</sub> 1h<br>IC<sub>80</sub> 4h | -6.64<br>-6.64 | Dogan et al. 2015 |
| P06493 | CDK1 | High levels of CDK1 expression in patients with lung squamous cell carcinoma are associated with worse prognosis. | IC<sub>80</sub> 1h | -1.2 | Deng et al. 2021 |
| Q5EC54 | Heterogeneous nuclear ribonucleoprotein K | hnRNP K plays a critical role in lung cancer progression by inhibiting the p53/p21/cyclin D1-dependent signaling pathway. | IC<sub>50</sub> 1h<br>IC<sub>80</sub> 1h<br>IC<sub>50</sub> 4h<br>IC<sub>80</sub> 4h | -6.64<br>-6.64<br>-6.64<br>-6.64 | Li et al. 2022 |
| A0A6Q8PGN6 | Calpastatin | The calpain–calpastatin system plays a pivotal role at various developmental stages, in diverse tissues, and in numerous physiological and pathological processes. | IC<sub>50</sub> 1h<br>IC<sub>80</sub> 1h<br>IC<sub>80</sub> 4h | -1.98<br>-1.27<br>-1.4 | Nian and Ma 2021 |
| B2RCM3 | Calpain-2 catalytic subunit | Calpain-2 expression was found to be up-regulated in NSCLC cell lines and tissues. | IC<sub>80</sub> 1h<br>IC<sub>50</sub> 4h<br>IC<sub>80</sub> 4h | -6.64<br>-6.64<br>-6.64 | Xu et al. 2019 |
| P37802 | Transgelin-2 | Comparative proteomics and immunohistochemistry revealed the upregulation of transgelin-2 in tumor-derived lung cancer endothelial cells in lung cancer patients. | IC<sub>50</sub> 1h<br>IC<sub>80</sub> 1h<br>IC<sub>80</sub> 4h | -1.87<br>-1.6<br>-1.63 | Jin et al. 2019 |
| Q16658 | Fascin | Fascin was found to be highly expressed in NSCLC tissues. Furthermore, high fascin-1 expression in NSCLC was significantly associated with lymph node metastasis. IC80 1h | IC<sub>80</sub> 1h<br>IC<sub>50</sub> 4h<br>IC<sub>80</sub> 4h | -1.43<br>-1.48<br>-2.15 | Ling et al. 2015 |
| A0A140VK08 | Calretinin | Calretinin-positive lung tumors exhibit similarities to those observed in smokers and individuals with advanced stages of disease. | IC<sub>50</sub> 1h<br>IC<sub>80</sub> 1h | -1.83<br>-1.21 | Matsuda et al. 2020 |
| Q9H773 | dCTP pyrophosphatase 1 | The expression of DCTPP1 was markedly upregulated in squamous NSCLC patients, accompanied by a notable increase in DCTPP1 nucleic expression. | IC<sub>80</sub> 1h<br>IC<sub>80</sub> 4h | -1.37<br>-1.28 | Zhang et al 2013 |

FC: factor change, t: time

## Discussion

Most of the anti-cancer drugs are designed to block or inhibit abnormally active proteins that drive tumor growth, angiogenesis, metastasis, and other cancer-linked processes. However, to fully understand the mechanism of action of the drugs it is crucial to characterize the proteome-wide scale in a dose- and time-depending setting. In line with this, the newly established proteomic technology (DecryptM) permits to generate large-scale biology data assessing cellular mechanism of action in a dose- and time-resolved fashion (Zecha et al. 2023). The implementation of this research approach in drug-regulated proteomics studies allow a better understanding of the pharmacological mechanism of anticancer drugs as well as the identification of proteins suitable for drug response biomarkers. In line with this reasoning, in this work we have interrogated the temporal proteome regulation by the clinical-grade CIGB-300 anticancer peptide on lung squamous NCI-H226 cells at two dose levels which produce 50% and 80% of cell death, respectively.

Globally, our data shown that down-regulation prevailed over up-regulation of proteins after treating NCI-H226 with CIGB-300 IC_50_ and IC_80_ during 1 or 4 hours. Interestingly, cells treated with IC_80_ exhibited higher number of down-regulated proteins compared to those treated with IC_50_ treatment, 577 *versus* 305 after 1 hour and 493 *versus* 287 after 4 hours of incubation. This inhibitory dose-dependent effect of CIGB-300 is very consistent with the dose-effect observed in terms of cell death burden which was higher at IC_80_. However, only marginal effect was observed in the up-regulated protein levels which did not exhibit a clear dose-dependent effect between IC_50_ and IC_80_. Although, both down- and up-regulated proteins can be involved on the CIGB-300 cell death burden, our data suggest that those down-regulated proteins could play preponderant role on the cytotoxic effect of CIGB-300. Furthermore, in the proteomic profile regulated at 1h and 4h, a time-dependent decrease in the number of modulated proteins was observed at both peptide doses, which was more evident in the subset of up-regulated proteins.

The functional enrichment analysis of the CIGB-300-regulated proteome in NCI-H226 cells illustrates that a dose-dependent effect characterizes the cellular response to both peptide concentrations. In general, the biological process and pathways modulated by CIGB-300 IC_50_ are also enriched in the proteomic profile modulated by the higher peptide dose. There is a core of biological processes related to small molecule biosynthesis, aldehyde metabolism and metabolism of nucleobases that are modulated irrespectively to the peptide dose or the treatment time. However, several functional annotations are only over-represented in NCI-H226 cells treated with CIGB-300 IC_80_. For instance, proteins related to drug ADME (Absorption, Distribution, Metabolism and Excretion) properties were over-represented in the proteomic profile after incubation with CIGB-300 IC_80_. The modulation of drug-related metabolic pathways may be associated with the cytotoxicity of higher doses of CIGB-300. Furthermore, the oncogenic signature NFE2L2.V2, which includes down-regulated genes after the inhibition of the transcription factor NFE2L2/NRF2 (Kim et al. 2016) [**Error! Bookmark not defined.**] was exclusively over-represented in the proteome of NCI-H226 cells treated with CIGB-300 IC_80_. The transcription factor NRF2 is frequently over-activated in many cancers including lung cancer (Lin et al. 2023), induces a cytoprotective response to oxidative stress and also activates the expression of genes involved in drug detoxification, inflammation and metabolism (He et al. 2020). Importantly, CK2 regulates the NRF2-mediated cellular antioxidant response (Pi et al. 2007). These results suggest that CIGB-300 could affect the NRF2 - mediated transcriptional events in H226-treated cells at the highest dose level. In line with such result, proteins related to glutathione metabolism were modulated in response to CIGB-300 IC_80_ treatment.

In our work, we also explored whether CIGB-300 dose and time of treatment did modulate individual proteins. To do that we were focused on 13 up-regulated proteins previously reported to be associated to NSCLC and squamous NSCLC. As revealed by the fold-change values respect to control, data from Table 1 indicated that treatment with CIGB-300 did affect the levels of all the proteins in two different ways, those modulated at IC_50_ and IC_80_ (LDBH, PIG3, hnRNPK, Calpastatin, Calpain-2, Transgelin-2, Fascin, Calretinin) and those modulated only at IC_80_ (Fructose-Biphosphate Aldolase, LDHA, MAPK, CDK1, dCTP-Pyrophosphate-1).

In tumor environment, glucose is limited, thus cancer cells resorts to nucleoside catabolism to obtain ribose-5-P which is then incorporated to the pentose phosphate pathway to generate glycolysis intermediates and therefore metabolic energy (Strefeler et al. 2024). Fructose-bisphosphate aldolase (ALDO) was down-regulated at IC_80_ dose at 1 and 4 h of incubation time. ALDO, which is involved in both glycolysis and gluconeogenesis, was associated in functional analysis to nucleobases-small containing molecules metabolic processes. The role of aldolases in the metastasis of different types of cancer has been confirmed by multiple studies which show that their action goes beyond glucose metabolism, but also participates in other pathways such as Wnt/β-catenin, EGFR/MAPK, Akt and HIF-1α ( Tang and Cui 2024).

Another enzyme related to energetic metabolism and highly expressed in tumor cells is Lactate Dehydrogenase (LDH A and B). The fermentation of glucose to lactate in cancer cells occurs even in the presence of sufficient oxygen to sustain mitochondrial oxidative phosphorylation with complete catabolism of glucose and maximizing ATP production (Warburg Effect, Liberti and Locasale 2016). For the other hand, some studies have indicated that there is a correlation between the presence of LDHB and the increased proliferation of lung adenocarcinoma (McCleland et al. 2013). LDH inhibition as a potential therapeutic strategy in cancer treatment was first proposed in the 1960s. In this work, both enzymes were inhibited by CIGB-300 IC80 at 1 and 4 h, and LDHB was also inhibited by IC50 at 4 h. Considering that CIGB-300 is a CK2 inhibitor, these results agree with previous studies that showed that protein level of LDHA and LDHB, was down-regulated in CK2α-silenced cells compared with those in control cells. Our group previously reported inhibition of LDH A and B by CIGB-300 peptide in NSCLC with a lower dose and 45 minutes of incubation time (Rodríguez-Ulloa et al. 2010).

Proteomics results showed a down-regulation of Quinone oxidoreductase PIG3 in all experimental conditions. This protein is associated with metabolism of nucleotides, which were also overrepresented at both CIGB-300 treatment times. This protein is involved in the response to cell damage and apoptosis, and previous studies have described it as a controversial enzyme regarding its role in the cancer phenotype. In NSCLC adenocarcinoma, it has been associated with cell progression and metastasis (Gu et al. 2018).

Another key protein involves in cancer cell proliferation, migration and survival that was inhibited by IC_80_ dose of CIGB-300 at 1h and 4h was the mitogen-activated protein kinase (MAPK1, MAP2K2, and MAP2K4), MAP2K2 and MAP2K4 were down-regulated by IC_50_ dose of CIGB-300 at 1h and 4h. These proteins are involved in MAP kinase signal transduction pathway, associated to nucleobases-small containing molecules metabolic processes and over-represented in the proteomic profile differentially modulated by IC_80_ at 1h and 4h in Functional enrichment analysis. Mitogen-activated protein kinase (MAPK or MEK) pathway modulates tumor cell survival and proliferation in non-small cell lung cancer. Recent data of MEK-inhibitor-therapy based Combination Strategies in NSCLC was reported (Goldman et al. 2012). Thus, CIGB-300 could be potential candidate to use in combination strategies for MEK inhibition in squamous NSCLC.

Heterogeneous nuclear ribonucleoprotein K was found to be selectively down-regulated in cells treated with both, IC_50_ and IC_80_ doses at 1 and 4 h. Similar result was previously reported, where CIGB-300 inhibits hnRNPK in non-small cell Lung cancer (Rodríguez-Ulloa et al. 2010). Heterogeneous nuclear ribonucleoproteins (hnRNP) K play an important role in the p53-triggered DNA damage response, acting as a cofactor for p53 in response to DNA damage (Pelisch et al. 2012).

Metabolism of nucleotide, one of the most enriched processes showed the presence of the enzyme dCTP pyrophosphatase 1 (DCTPP1). This protein was down-regulated in samples treated with IC_80_ dose of CIGB-300 peptide. According to Gene Ontology annotation its function is to convert deoxycytidine triphosphate (dCTP) to deoxycytidine monophosphate (dCMP) and inorganic pyrophosphate and it has been demonstrated to be involved in numerous diseases, including several types of cancer. A study about breast cancer proved that high intracellular concentrations of the enzyme were related to a poor prognosis of patients and that its tumorigenic effect depended on the DNA repair signaling pathway (Niu et al. 2021). Additionally, the expression of DCTPP1 was markedly up-regulated in squamous NSCLC patients (Zhang et al. 2013).

Transgelin-2, Fascin, Calpain–calpastatin system and Calretinin were found to be selectively down-regulated in NCI-H226 cells treated with both IC50 and IC80 doses of CIGB–300. These and all proteins here discussed are biomarkers or potential markers of NSCLC and squamous NSCLC, some of them are involved in a selective pathway of tumoral cells, thereby supporting the hypothesis of the therapeutic potential of CIGB-300 anticancer treatment for this kind of tumor.

The fact that we have found proteins associated with the tumor biology and the prognosis of lung cancer patients, allows us to suppose that such proteins could also be candidates for pharmacodynamic biomarkers of CIGB-300 in lung cancer. This hypothesis deserves to be confirmed in the context of clinical trials of CIGB-300 in patients with lung cancer.

Although this quantitative proteomics strategy in our work does not provide direct information on the phosphorylation status of proteins, we investigated a possible modulatory effect of CIGB-300 on the abundance of CK2 substrates like previously found with this CK2 inhibitor in comparative proteomics studies. Interestingly, the results showed that CK2 substrates modulated by CIGB-300 interact with differentially modulated proteins that support the survival and proliferative capacity of NCI-H226 cells. In addition, the interactome around CK2 substrates is more densely interconnected with the proteome regulated by CIGB-300 IC_80_. At both peptide concentrations, CK2 substrates modulated by CIGB-300 interact with proteins related to glycolysis, the pentose phosphate pathway, ubiquitination and cell death. The functional association between CK2 substrates and proteins related to the MAPK cascade, the cell cycle and the detoxification process was only evident at the higher peptide dose.

Overall, our findings describe for the first time a quantitative proteomics analysis of CIGB-300 in lung squamous cancer cells in a dose- and time dependent setting. By increasing the dose of CIGB-300 entails the differential appearance of processes and signalling cascades mainly linked to the anticellular effect of CIGB-300 while a preselected group of proteins linked to the biology of the disease and/or the prognosis of the patients were significantly inhibited irrespectively of drug dose and time of incubation.

## Conclusions

For the first time, a quantitative proteomic analysis was performed in dose- and time dependent fashion with CIGB-300 in squamous lung cancer cells. A number of proteins were differentially modulated by CIGB-300 and most of them were found to be down-regulated, indicating a cytotoxic and pro-apoptotic effect of the peptide, particularly pronounced in selective pathways of tumoral cells such as: aerobic glycolysis (Warburg Effect) and pentose phosphate pathways. The biological processes that were more represented among the modulated proteins were those related to small molecule biosynthetic process, pyridine-containing compound metabolic process and nucleobase-containing small molecule metabolism. LUAD and LUSC previously described biomarkers were negatively modulated by treatment with both CIGB-300 doses. These proteins, which are associated with metabolism and cell survival, reinforce the hypothesis of the therapeutic potential of CIGB-300 as anticancer treatment for squamous non-small cell lung cancer and merits be validated as CIGB-300 response biomarkers in NSCLC patients.

## Supporting information

Protein Identification

## Acknowledgements

The authors would like to thank to Ke Yang and Changyuan Tan for administrative and logistic supporting of this work.

## Availability of data and materials

The authors disclose that all materials and data generated in this work is provided in the submitted article and can be shared upon request.

## Authors’ contributions

YP and SEP conceived the original research idea for the project. YP is the principal investigator of the project. LGH, ARU and SEP wrote the manuscript and data analysis. WL contributed to the conceptualization and overall supervision. LD and YY conducted the experimentation in this work. Both LJG and VBL co-supervised the work and helped with interpretation of data.

## Ethics approval and consent to participate

We declare that all of the authors approved this manuscript and expressed their consent and willingness to participate as authors in this work.

## References

Ashburner, M., Ball, C. A., Blake, J. A., Botstein, D., Butler, H., Cherry, J. M. et al. (2000) Gene ontology: tool for the unification of biology. The Gene Ontology Consortium. Nat Genet 25, 25–29.

Batista-Albuerne N., González-Méndez L., García-García I., Fernández-Sánchez E., García-Diegues R., et al. (2018) Phase I Study of CIGB-300 Administered Intravenously in Patients with Relapsed/Refractory Solid Tumors. J Med Oncol. 1:4.

Borgo, C., D’Amore, C., Sarno, S., Salvi, M., & Ruzzene, M. (2021). Protein kinase CK2: a potential therapeutic target for diverse human diseases. Signal transduction and targeted therapy, 6(1): 183.

Bray F., Laversanne M., Sung H., et al. (2024). Global cancer statistics 2022: GLOBOCAN estimates of incidence and mortality worldwide for 36 cancers in 185 countries. CA Cancer J Clin. 74(3): 229–263.

Deng H., Gao Y., Trappetti V., Hertig D., Karatkevich D., Losmanova T., Urzi C., Ge H., Geest G.A., Bruggmann R., Djonov V., Nuoffer J.M., Vermathen P., Zamboni N., Riether C., Ochsenbein A., Peng R.W., Kocher G.J., Schmid R.A., Dorn P., Marti T.M. (2022). Targeting lactate dehydrogenase B-dependent mitochondrial metabolism affects tumor initiating cells and inhibits tumorigenesis of non-small cell lung cancer by inducing mtDNA damage. Cell Mol Life Sci. 79:445. doi: 10.1007/s00018-022-04453-5.

Deng, H., Hang, Q., Shen, D., Ying, H., Zhang, Y., Qian, X., & Chen, M. (2021). High Expression Levels of CDK1 and CDC20 in Patients With Lung Squamous Cell Carcinoma are Associated With Worse Prognosis. Frontiers in Molecular Biosciences, 8: 653805.

Dogan Turacli, A.C. and Ozkan, A. Ekmekci. (2015). The comparison between dual inhibition of mTOR with MAPK and PI3K signaling pathways in KRAS mutant NSCLC cell lines Tumor Biol., 36: 9339–9345.

Du S., Guan Z., Hao L., Song Y, Wang L, Gong L, et al. (2014) Fructose-Bisphosphate Aldolase A Is a Potential Metastasis-Associated Marker of Lung Squamous Cell Carcinoma and Promotes Lung Cell Tumorigenesis and Migration. PLoS ONE 9: e85804. 10.1371/journal.pone.0085804

Fiume, L. (1960). Inhibition of Aerobic Glycolysis in Yoshida Ascites Hepatoma by Tartronic Acid. Nature 187: 792–793. doi.org/10.1038/187792a0.

Gillespie, M., Jassal, B., Stephan, R., Milacic, M., Rothfels, K., Senff-Ribeiro, A. et al. (2022) The reactome pathway knowledgebase 2022. Nucleic Acids Res 50: D687–D692.

Goldman J.W. and Garon E.B. (2012). Targeting MEK for the treatment of non-small-cell lung cancer. J Thorac Oncol. 16:S377–8. doi: 10.1097/JTO.0b013e31826df0bc.

Gu, M. M., Gao, D., Yao, P. A., Yu, L., Yang, X. D., Xing, C. G. et al. (2018). p53-inducible gene 3 promotes cell migration and invasion by activating the FAK/Src pathway in lung adenocarcinoma. Cancer science, 109: 3783–3793.

He F., Ru X., Wen T. (2020). NRF2, a Transcription Factor for Stress Response and Beyond. Int J Mol Sci. 21:4777.

Herbst, R. S., Morgensztern, D. and Boshoff, C. (2018). The biology and management of non-small cell lung cancer. Nature 553: 446–454.

Hornbeck P.V., Kornhauser J.M., Latham V., Murray B., Nandhikonda V., Nord A., Skrzypek E., Wheeler T., Zhang B., Gnad F. (2019). 15 years of PhosphoSitePlus®: integrating post-translationally modified sites, disease variants and isoforms. Nucleic Acids Res. 47:D433–D441. doi: 10.1093/nar/gky1159.

Jin, H., Cheng, X., Pei, Y., Fu, J., Lyu, Z., Peng, H., Yao, Q., Jiang, Y., Luo, L., & Zhuo, H. (2016). Identification and verification of transgelin-2 as a potential biomarker of tumor-derived lung-cancer endothelial cells by comparative proteomics. Journal of proteomics 136: 77–88.

Kanehisa, M. and Goto, S. (2000) KEGG: kyoto encyclopedia of genes and genomes. Nucleic Acids Res 28: 27–30.

Kim J.W., Botvinnik O.B., Abudayyeh O., Birger C., Rosenbluh J., Shrestha Y., Abazeed M.E., Hammerman P.S., DiCara D., Konieczkowski D.J., Johannessen C.M., Liberzon A., Alizad-Rahvar A.R., Alexe G., Aguirre A., Ghandi M., Greulich H., Vazquez F., Weir B.A., Van Allen E.M., Tsherniak A., Shao D.D., Zack T.I., Noble M., Getz G., Beroukhim R., Garraway L.A., Ardakani M., Romualdi C., Sales G., Barbie D.A., Boehm J.S., Hahn W.C., Mesirov J.P., Tamayo P. (2016). Characterizing genomic alterations in cancer by complementary functional associations. Nat Biotechnol. 34:539–46.

Li M., Li S., Liu B., Gu M.M., Zou S., Xiao B.B., Yu L., Ding W.Q., Zhou P.K., Zhou J., Shang Z.F. (2017). PIG3 promotes NSCLC cell mitotic progression and is associated with poor prognosis of NSCLC patients. J Exp Clin Cancer Res. 36:39. doi: 10.1186/s13046-017-0508-2.

Li M., Yang X., Zhang G., Wang L., Zhu Z., Zhang W., Huang H., Gao R. (2022). Heterogeneous nuclear ribonucleoprotein K promotes the progression of lung cancer by inhibiting the p53-dependent signaling pathway. Thorac Cancer. 13:1311–1321. doi: 10.1111/1759-7714.14387.

Liao, R. G., Watanabe, H., Meyerson, M., & Hammerman, P. S. (2012). Targeted therapy for squamous cell lung cancer. Lung cancer management, 1: 293–300.

Liberti M.V. and Locasale J.W. (2016). The Warburg Effect: How Does it Benefit Cancer Cells? Trends Biochem Sci. 41:211–218. doi: 10.1016/j.tibs.2015.12.001. Erratum in: Trends Biochem Sci. 41:287.

Liberzon A., Birger C., Thorvaldsdóttir H., Ghandi M., Mesirov J.P., Tamayo P. (2015). The Molecular Signatures Database (MSigDB) hallmark gene set collection. Cell Syst. 23:417–425. doi: 10.1016/j.cels.2015.12.004.

Lin L., Wu Q., Lu F., Lei J., Zhou Y., Liu Y., Zhu N., Yu Y., Ning Z., She T., Hu M. (2023). Nrf2 signaling pathway: current status and potential therapeutic targetable role in human cancers. Front Oncol. 22:1184079.

Ling X.L., Zhang T., Hou X.M., Zhao D. (2015). Clinicopathological significance of fascin-1 expression in patients with non-small cell lung cancer. Onco Targets Ther. Jun 30:1589–95. doi: 10.2147/OTT.S84308.

Matsuda M., Ninomiya H., Wakejima R., Inamura K., Okumura S., Mun M., Kitagawa M., Ishikawa Y. (2020) Calretinin-expressing lung adenocarcinoma: Distinct characteristics of advanced stages, smoker-type features, and rare expression of other mesothelial markers are useful to differentiate epithelioid mesothelioma. Pathology, Research and Practice. 216:152817. DOI: 10.1016/j.prp.2020.152817.

McCleland, M.L., Adler, A.S., Deming, L., Cosino, E., Lee, L., Blackwood, E.M., Solon, M., Tao, J., Li, L., Shames, D., et al. (2013). Lactate dehydrogenase B is required for the growth of KRAS-dependent lung adenocarcinomas. Clin. Cancer Res. 19: 773–784.

Nian H. and Ma B. (2021). Calpain-calpastatin system and cancer progression. Biol Rev Camb Philos Soc. 96:961–975. doi: 10.1111/brv.12686.

Niu, M., Shan, M., Liu, Y., Song, Y., Han, J.-g., Sun, S., Zhang, G.Q. (2021). DCTPP1, an oncogene regulated by miR-378a-3p, promotes proliferation of breast cancer via DNA repair signaling pathway. Frontiers in oncology, 11: 641931. doi: 10.3389/fonc.2021.641931.

Niu, Z., Jin, R., Zhang, Y., & Li, H. (2022). Signaling pathways and targeted therapies in lung squamous cell carcinoma: mechanisms and clinical trials. Signal Transduction and Targeted Therapy, 7: 353–353. doi.org/10.1038/s41392-022-01200-x.

Pelisch F., Pozzi B., Risso G., Muñoz M. J. and Srebrow A. (2012) DNA damage-induced heterogeneous nuclear ribonucleoprotein K sumoylation regulates p53 transcriptional activation J. Biol. Chem. 287: 30789–99.

Perea, S. E., Reyes, O., Puchades, Y., Mendoza, O., Vispo, N. S., Torrens, I., Santos, A., Silva, R., Acevedo, B., López, E. et al. (2004). Antitumor Effect of a Novel Proapoptotic Peptide that Impairs the Phosphorylation by the Protein Kinase 2 (Casein Kinase 2). Cancer Res. 64: 7127–7129. doi:10.1158/0008-5472.

Perea, S. E., Reyes, O., Baladron, I., Perera, Y., Farina, H., Gil, J., Rodríguez, A., Bacardí, D., Marcelo, J.L., Cosme, K. et al. (2008). CIGB-300, a Novel Proapoptotic Peptide that Impairs the CK2 Phosphorylation and Exhibits Anticancer Properties Both In Vitro and In Vivo. Mol. Cel Biochem 316: 163–167. doi:10.1007/s11010-008-9814-5.

Perea, S. E., Baladrón, I., Valenzuela, C. and Perera, Y. CIGB-300: a peptide-based drug that impairs the protein kinase CK2-mediated phosphorylation. Semin. Oncol. 45: 58–67.

Perera, Y., Farina, H.G.; Gil, J.; Rodriguez, A.; Benavent, F.; Castellanos, L.; Gómez, R.E.; Acevedo, B.E.; Alonso, D.F.; Perea, S.E. et al. (2009). Anticancer peptide CIGB-300 binds to nucleophosmin/B23, impairs its CK2-mediated phosphorylation, and leads to apoptosis through its nucleolar disassembly activity. Mol. Cancer Ther. 8: 1189–1196.

Perera, Y., Ramos, Y., Padrón, G., Caballero, E., Guirola, O., Caligiuri, L.G., Lorenzo, N., Gottardo, F., Farina, H.G., Filhol, O.;, et al. (2020). CIGB-300 anticancer peptide regulates the protein kinase CK2-dependent phosphoproteome. Mol. Cell. Biochem. 470: 63–75.

Perera, Y., Pedroso, S., Borras-Hidalgo, O., Vázquez, D. M., Miranda, J., Villareal, A., Falcón V., Cruz L. D., Farinas H. G., Perea S. E. (2015). Pharmacologic Inhibition of the CK2-Mediated Phosphorylation of B23/NPM in Cancer Cells Selectively Modulates Genes Related to Protein Synthesis, Energetic Metabolism, and Ribosomal Biogenesis. Mol. Cel Biochem 404: 103–112. doi:10.1007/s11010-015-2370-x.

Pi J., Bai Y., Reece J.M., Williams J., Liu D., Freeman M.L., Fahl W.E., Shugar D., Liu J., Qu W., Collins S., Waalkes M.P. (2007). Molecular mechanism of human Nrf2 activation and degradation: role of sequential phosphorylation by protein kinase CK2. Free Radic Biol Med. 42:1797–806.

Plotnikov A., Chuderland D., Karamansha Y., Livnah O., Seger R. (2019). Nuclear ERK Translocation is Mediated by Protein Kinase CK2 and Accelerated by Autophosphorylation. Cellular Physiology and Biochemistry: International Journal of Experimental Cellular Physiology, Biochemistry, and Pharmacology. 53:366–387. DOI: 10.33594/000000144.

Rodríguez-Ulloa, A., Ramos, Y., Gil, J., Perera, Y., Castellanos-Serra, L.R., García, Y., Betancourt, L.H., Besada, V., González, L.J., Fernández-de-Cossio, J., Sánchez, A., Serrano, J.M., Farina, H.G., Alonso, D.F., Acevedo, B.E., Padrón, G., Musacchio, A., & Perea, S.E. (2010). Proteomic profile regulated by the anticancer peptide CIGB-300 in non-small cell lung cancer (NSCLC) cells. Journal of Proteome Research 9: 5473–83.

Shannon P., Markiel A., Ozier O., Baliga N.S., Wang J.T., Ramage D., Amin N., Schwikowski B., Ideker T. Cytoscape: a software environment for integrated models of biomolecular interaction networks. (2003). Genome Res. 13:2498–504. doi: 10.1101/gr.1239303.

Strefeler, A., Blanco-Fernandez, J., and Jourdain, A. A. (2024). Nucleosides are overlooked fuels in central carbon metabolism. Trends in Endocrinology & Metabolism. 35: 290–299.

Szklarczyk D., Gable A.L., Lyon D., Junge A., Wyder S., Huerta-Cepas J., Simonovic M., Doncheva N. T., Morris J. H., Bork P. et al. (2019). STRING v11: protein–protein association networks with increased coverage, supporting functional discovery in genome-wide experimental datasets, Nucleic Acids Research 47: D607–D613, 10.1093/nar/gky1131.

Tang, F., and Cui, Q. (2024). Diverse roles of aldolase enzymes in cancer development, drug resistance and therapeutic approaches as moonlighting enzymes. Medical Oncology 41: 224.

Wiśniewski J.R., Zougman A., Nagaraj N., Mann M. (2009). Universal sample preparation method for proteome analysis. Nat Methods. 6:359–62. doi: 10.1038/nmeth.1322.

Xu F., Gu J., Lu C., Mao W., Wang L., Zhu Q., Liu Z., Chu Y., Liu R., Ge D. (2019). Calpain-2 Enhances Non-Small Cell Lung Cancer Progression and Chemoresistance to Paclitaxel via EGFR-pAKT Pathway. Int J Biol Sci. 15:127–137. doi: 10.7150/ijbs.28834.

Yang Y., Chong Y., Chen M., Dai, W., Zhou, X., Ji Y., Qiu G., Du X. et al. (2021). Targeting lactate dehydrogenase a improves radiotherapy efficacy in non-small cell lung cancer: from bedside to bench. J Transl Med 19: 170. 10.1186/s12967-021-02825-2.

Zecha J., Bayer F.P., Wiechmann S., Woortman J., Berner N., Müller J., Schneider A., Kramer K., Abril-Gil M., Hopf T. et al. (2023). Decrypting drug actions and protein modifications by dose- and time-resolved proteomics. Science 380: 93–101. DOI: 10.1126/science.ade3925.

Zhang X., Yang X., Yang C., Li P., Yuan W., Deng X., Cheng Y., Li P., Yang H., Tao J., Lu Q. (2016). Targeting protein kinase CK2 suppresses bladder cancer cell survival via the glucose metabolic pathway. Oncotarget. 7: 87361–87372. doi: 10.18632/oncotarget.13571.

Zhang Y., Ye W.Y., Wang J.Q., Wang S.J., Ji P., Zhou G.Y., Zhao G.P., Ge H.L., Wang Y. (2013). dCTP pyrophosphohydrase exhibits nucleic accumulation in multiple carcinomas. Eur J Histochem. 57:e29. doi: 10.4081/ejh.2013.e29.

Zheng Y., Qin H., Frank S.J., Deng L., Litchfield D.W., Tefferi A., Pardanani A., Lin F.T., Li J., Sha B., Benveniste E.N. (2011) A CK2-dependent mechanism for activation of the JAK-STAT signaling pathway. Blood 118:156–66. doi: 10.1182/blood-2010-01-266320.

Zhou, Y., Zhou, B., Pache, L., Chang, M., Khodabakhshi, A. H., Tanaseichuk, O. et al. (2019) Metascape provides a biologist-oriented resource for the analysis of systems-level datasets. Nat Commun 10: 1523.

